# Detection and Purification of Lewy Pathology from Formalin Fixed Primary Human Tissue Using Biotinylation by Antigen Recognition

**DOI:** 10.1101/2020.11.11.378752

**Authors:** BA Killinger, L Marshall, D Chatterjee, Y Chu, JH Kordower

**Affiliations:** Department of Neurological Sciences, Rush University Medical Center, Chicago Illinois 60612; Center for Neurodegenerative Science, Van Andel Research Institute, Grand Rapids MI 49503

**Keywords:** Prion disease, Neurodegeneration, Proximity labeling, In-situ, Mass spectrometry, Exosomes

## Abstract

The intracellular misfolding and accumulation of alpha-synuclein into structures collectively called Lewy pathology is a central phenomenon for the pathogenesis of Parkinson’s disease (PD), Dementia with Lewy Bodies (DLB), and Multiple System Atrophy. Understanding the molecular architecture of Lewy pathology is crucial for understanding disease origins and progression. Here we developed a method to label, extract, and purify molecules from Lewy pathology of formalin fixed PD and DLB brain for blotting and mass spectrometry analysis. Using the biotinylation antibody recognition (BAR) technique, we labeled phosphoserine 129 alpha-synuclein positive pathology and associated molecules with biotin. Formalin crosslinks were then reversed, protein extracted, and pathology associated molecules isolated with streptavidin beads. Results showed superior immunohistochemical staining of Lewy pathology following the BAR protocol when compared to standard avidin biotin complex (ABC) based detection. The enhanced staining was particularly apparent for fibers of the medial forebrain bundle and punctate pathology within the striatum and cortex, which otherwise were weakly labeled or not detected. Subsequent immunoblotting BAR-labeled Lewy pathology extracts revealed the presence of high molecular weight alpha-synuclein, ubiquitin protein conjugates, and phosphoserine 129 alpha-synuclein. Mass spectrometry analysis of BAR-labeled Lewy pathology extracts from PD and DLB patients identified 815 proteins with significant enrichment for many pathways. Notably the most significant KEGG pathway was Parkinson’s disease (FDR = 2.48 × 10^−26^) and GO Cellular compartment was extracellular exosomes (GO Cellular Compartment; FDR = 2.66× 10^−34^). We used enrichment data to create a functional map of Lewy Pathology from primary disease tissues, which implicated Vesicle Trafficking as the primary disease associated pathway in DLB and PD. In summary, this protocol can be used to enrich for Lewy pathology from formalin fixed human primary tissues, which allows the determination of molecular signatures of Lewy pathology. This technique has broad potential to help understand the phenomenon of Lewy pathology in primary human tissue and animal models.

## Introduction

Parkinson’s disease (PD) and Dementia with Lewy Bodies (DLB) are progressive neurodegenerative diseases characterized by intracellular inclusions containing aggregated alpha-synuclein referred to as Lewy bodies or Lewy neurites, depending on the cellular compartment in which it is observed (i.e. cell bodies and nerve fibers, respectively). Although it is clear that alpha-synuclein is involved in the disease process, central questions remain regarding the molecular origins and progression of Lewy pathology. A diverse range of cellular pathways have been implicated in alpha-synuclein disease processes including dysfunctional synaptic neurotransmission and vesicular trafficking, mitochondrial dysfunction, neuroinflammation, calcium signaling, glycolysis, autophagy lysosomal dysfunction, and MAPK signaling pathways^1^. However, much of the evidence implicating these cellular processes have been derived from cell and animal models, which may or may not closely recapitulate the human disease processes.

Alpha-synuclein phosphorylated at serine 129 (PSER129) is a marker of Lewy pathology. PSER129 is enriched in Lewy pathology with only trace quantities (<5% of total alpha-synuclein pool) of endogenous alpha-synuclein being phosphorylated^2^. Evidence suggests that PSER129 may aggregate more rapidly and PSER129 may have a role in regulating alpha-synuclein turnover^3^. Evidence suggests that PSER129 accumulates in the brain as PD disease progresses^4,5^. Although it is not clear if PSER129 occurs prior to, during, or following inclusion formation^6^, it likely occurs once alpha-synuclein aggregation has initiated. Several antibodies have been developed to detect PSER129, including the commonly used EP1536Y antibody (Abcam), a monoclonal antibody against PSER129 that has high specificity for Lewy pathology^7^ and has been used extensively for immunohistochemically (IHC) labeling of Lewy pathology.

Purification of Lewy pathology from primary tissue typically requires fresh (i.e. unfixed) tissue, cellular disruption, and extensive fractionation, which disturbs both cellular structures, and presumably, Lewy body composition. Formalin fixed samples preserve cellular morphology and preserve Lewy pathology in its native state; however, extraction of proteins, protein complexes, and organelles from formalin fixed tissues is challenging, and often impractical. Recently, a technique referred to as biotinylation by antigen recognition (BAR) was successfully used to label cellular components within close proximity to an antigen in fixed primary tissues for enrichment and downstream mass spectrometry analysis^8–11^. The BAR technique [also called selective proteomic proximity labeling using tyramide (SPPLAT) or enzyme-mediated activation of radical sources (EMARS)], labels a target antigen by peroxidase-catalyzed deposition of biotin moieties onto proximal molecules. BAR is particularly well suited for determining protein interactions and subcellular localization of insoluble cellular components^8^. Lewy pathology are insoluble cellular inclusions and, therefore, the BAR technique holds promise to study these structures. Recent demonstrations of the BAR technique, and its utility for proteomic analysis, gave us the impetus to attempt and purify Lewy pathology from fixed primary human tissues.

Here we applied a modified BAR technique referred to as ABC-BAR-PSER129 in formalin-fixed human brain samples from individuals diagnosed with PD and DLB. Several important results were obtained: 1) ABC-BAR-PSER129 technique used for labeling provided superior immunohistochemical sensitivity and comprehensiveness of alpha-synuclein pathology labeling when compared to standard detection methods; 2) purified ABC-BAR-PSER129 pathology was amenable to both western blot and mass spectrometry analysis; 3) fractions obtained with this protocol are significantly enriched for known PD pathways including vesicle trafficking; 4) extracellular exosomes were the most significantly enriched for go cellular compartment. Together, the ABC-BAR-PSER129 technique provides a practical way to detect and enrich for Lewy pathology in standard formalin fixed primary tissues.

## Methods

### Brain tissue preparation

Brains were immersion fixed in formalin for several days, equilibrated and cryoprotected in 30% sucrose solution, and 40-micron frozen sections were collected on a sliding knife microtome. Sections were selected from three PD and three DLB patients for immunostaining and purification (See table 1). Sections were stored in cryopreservation buffer (PBS pH 7.2, 30% sucrose, 30% ethylene glycol) at −20°C until processed. Tissue sections were in storage variable time before processing.

**Table 1.** Disease and demographic data for tissue used in study.

| Case ID | Diagnosis | Age (yrs) | Gender | PMI (h) | Disease duration (yrs) | UPDRS | H&Y |
| --- | --- | --- | --- | --- | --- | --- | --- |
| 1 | DLB | 87 | M | Unknown | 10 | 31 | 3 |
| 2 | DLB | 94 | F | 9 | Unknown | Unknown | Unknown |
| 3 | DLB<sup>1</sup> | 79 | M | 3 | 7 | 49 | 4 |
| 4 | PD<sup>1</sup> | 83 | M | 5 | 7 | 54 | 4 |
| 5 | PD | 72 | M | 10 | Unknown | 70 | 5 |
| 6 | PD | 81 | M | 5 | 13 | 45.5 | 4 |

<sup>1</sup>Sample analyzed by mass spectrometry. PMI = Post-mortem interval. UPDRS = Unified Parkinson's Disease Rating Scale. H&Y = Hoehn and Yahr scale.

### BAR labeling of formalin fixed floating sections

The workflow for the following studies are summarized in Figure 1. Floating sections were rinsed two times for 10 min with PBS, post-fixed in formalin for 30 min at room temperature and washed an additional two times with PBS. To quench endogenous peroxidase, sections were incubated for 30 min in PBS containing 3% H2O2 (Sigma). Following quenching, sections were briefly rinsed in PBS and then incubated in primary blocking buffer (1 × TBS pH 7.6, 0.5% triton X-100, 2% bovine serum albumin, 2% normal goat serum, 0.05% sodium azide) for 1h at room temperature. The PSER129 specific antibody EP1536Y (Abcam) was diluted in primary blocking buffer 1:50,000. Sections were then incubated with the primary antibody solution overnight (≥16h) at 4°C. For preabsorption studies, we incubated 1μl of EP1536Y and 20μl synthetic PSER129 peptide (Abcam) in 1mL blocking buffer overnight at 4°C. The preabsorbed mixture was then added to50mL of primary blocking buffer and incubated with tissue sections overnight at 4°C. For proteinase K digestion, prior to the fixation step sections were incubated in digestion buffer (1 × TBS pH 7.6, 0.5% triton X-100, 20 μg proteinase K / mL) for 10 min at 37°C, then post-fixed the sections for 105 minutes in formalin, and immunostaining protocol commenced.

**Figure 1.**
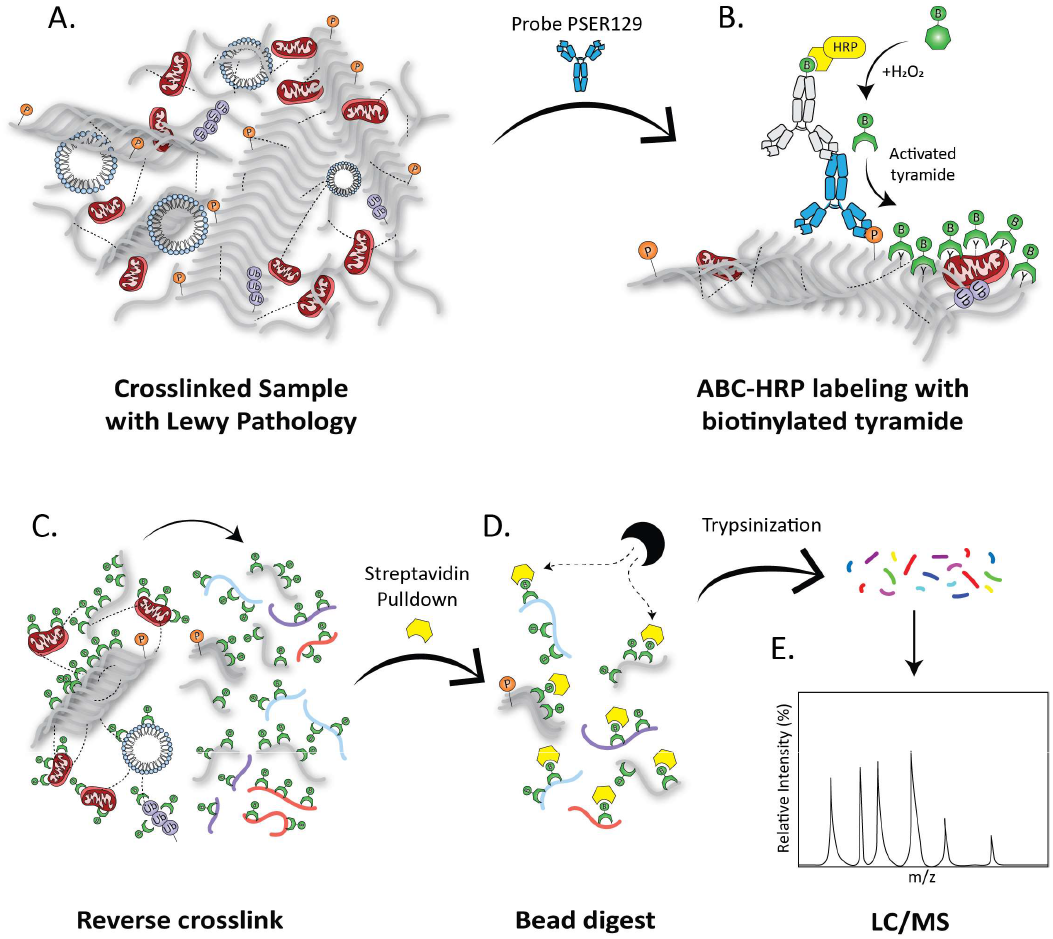
Strategy for BAR Labeling of Lewy Pathology. (A) Formalin fixed floating human brain sections were probed with anti-PSER129 antibody EP1536Y. (B) Using a biotinylated secondary antibody and avidin biotin complex, horse radish peroxidase (HRP) was then concentrated at the EP1536Y antigenic site. Biotin moieties were deposited near the antibody complex via peroxidase catalyzed oxidation of biotinyl tyramine. (C) Once Lewy pathology was labeled with biotin, formalin crosslinks were reversed. (D) The liberated biotin labeled Lewy pathology associated proteins could then be purified with streptavidin magnetic beads (E) and digested with trypsin for analysis via liquid chromatography mass spectrometry (LC/MS).

The next day sections were wash 6 × 10 min with wash buffer (1 × TBS, 0.05% triton X-100) and incubated with biotinylated antibody (Vector Laboratories) diluted 1:200 in secondary blocking buffer (1 × TBS pH 7.6, 0.05% triton X-100, 2% bovine serum albumin, 2% normal goat serum). Sections were then washed 3 × 10 min with wash buffer and incubated with ABC detection reagent (Vector Laboratories) for 1h at room temperature. Sections were then washed 2 × 10 min with wash buffer followed by 3 × 10 min in borate buffer (0.1M Sodium borate pH 8.5). To begin the BAR labeling reaction tissues were incubated with 100mL of borate buffer containing 2μl of stock biotinyl tyramide (Sigma, 5mg/mL dissolved in DMSO) and 10μl of 30% H202 (Sigma-Aldrich) for 30 min at room temperature. Sections were then washed 2 × 10 min with TBS and then 3 × 10 min with wash buffer. Once BAR labeled, tissue sections can be further processed or stored in wash buffer at 4°C.

### Detection of labeled processes

Once biotin labeled sections were washed, they could then be probed with ABC reagent as before to detect biotinylation of the tissue. Sections were developed using nickel-enhanced 3,3′-Diaminobenzidine (DAB) for 10 min. Following DAB development, sections were counterstained with methyl green, dehydrated in ethanol, cleared with xylene, and cover slipped with Cystoseal 60 mounting medium (ThermoFisher Scientific) for viewing under the microscope.

### Capture of biotinylated proteins

To remove biotin labeled antibody from tissues, sections were washed for 1h in antibody removal buffer (PBS, 2% SDS, 5% 2-Mercapitoethanol) similar to what has been previously described^12^. Sections were then rinsed with PBS 3 × 10 min. Sections were placed into 1.5 mL conical tube with 0.5mL of crosslink reversal buffer (500mM Tris-HCl pH 8.0, 150mM NaCl, 5% SDS, 2mM EDTA pH 8.0, and 2mM PMSF) and heated to 98°C for 30 min and 90°C for 2h on a heat block (ThermoFisher). Samples were mixed well and centrifuged at 22,000 × g for 30 min at room temperature. The supernatant was diluted to 3mL with TBST (1x TBS containing 1% triton x-100) and used for the capture protocol.

50μl of streptavidin magnetic beads (Thermofisher) were washed according to manufactures protocol and incubated with 3 mL of labeled tissue extract for 1 h at room temperature. Beads were isolated using a magnetic stand (Millipore) and washed 3 × 10 min with RIPA buffer (20mM tris-hcl pH 8.0, 150mM NaCl, 0.1% SDS, 1% Tx100) and then overnight with TBST (1x TBS containing 1% triton x-100). The lysate was retained following the pulldown for later characterization. Capture proteins were eluted by heating (98°C) the beads in 100 μl of SDS-PAGE sample buffer (ThermoFisher) for 10 min. Beads were collected using the magnetic stand and the eluent was retained for western blot. For mass spectrometry, beads were washed additionally 3 × 10 min with TBS to remove Triton X-100. Beads were collected and stored at −80°C until digestion for mass spectrometry.

### Western blot

5μl lysate and 25μl of bead eluent were resolved on 4-20% Bis-tris gels (ThermoFisher) and blotted onto activated polyvinylidene difluoride (PVDF) using wet transfer with settings at 120V for 1h. Following transfer membranes were rinsed with PBS and then fixed with 4% paraformaldehyde as previously described^13^. Membranes were then dried completely and reactivated with methanol just prior to use. Membranes were blocked in TBST containing 5% bovine serum albumin for 2 h at room temperature. Membranes were then incubated overnight at 4°C with blocking buffer containing one of the following antibodies; 1) anti-alpha-synuclein “SYN1” (BD Lifesciences) and “MJFR1” (Abcam), 2) anti-Ubiquitin-conjugates “UBCJ2” (Enzo Life Sciences), or 3) anti-PSER129 “EP1536Y” (Abcam). To determine enrichment of biotinylated proteins one membrane was incubated with prepared ABC reagent, diluted 1:10 in blocking buffer for 1h at room temperature (Vectorlabs). The next day, membranes were washed 3 × 10 min with TBST and incubated for 1 h with the appropriate secondary antibody (Cell Signaling Technologies) diluted in blocking buffer. Membranes were then wash 3 × 10 min with TBST and detected using chemiluminescence substrate (Bio-Rad).

### Mass spectrometry

ABC-BAR-PSER129 labeling was conducted on a putamen and midbrain section from two individuals, one diagnosed with PD and the other DLB (See table 1). Streptavidin beads containing the captured biotinylated proteins were washed and resuspended in a Tris/ Urea buffer. These were then reduced with dithiothreitol, alkylated with iodoacetamide, and digested with trypsin at 37°C overnight. This solution was subjected to solid phase extraction to concentrate the peptides and remove unwanted reagents followed by injection onto a Waters NanoAcquity HPLC equipped with a self-packed Aeris 3.6 μm C18 analytical column 0.075 mm by 20 cm, (Phenomenex). Peptides were eluted using standard reverse-phase gradients. The effluent from the column was analyzed using a Thermo Orbitrap Elite mass spectrometer (Nanospray configuration) operated in a data dependent manner for the 54 minutes. The resulting fragmentation spectra were correlated against the Refseq entries for homosapien using PEAKS Studio 10.5 (Bioinformatic Solutions). Proteins were retained for analysis that had at least one unique peptide identified and several assay contaminant proteins (i.e. streptavidin, bovine serum albumin, and trypsin) were removed from data prior to analysis.

### Data analysis

Pearson correlation analysis was conducted and all graphs created using GraphPad statistical software. Pathway enrichment was conducted using gProfiler^14^ using 10,075 proteins identified by shotgun mass spectrometry in bulk brain tissue as background^15^. Data presented was from an unordered query with a significance threshold of 0.05. Cytoscape (v.3.8.1), Enrichmentmap (v.3.3), AutoAnnotate (v.1.3.3) were used to generate pathway enrichment map16. Venny 2.117 was used to generate Venn diagrams.

## Results

First, we assessed the efficacy of biotin labeling using the ABC-BAR-PSER129 technique in the putamen and midbrain of an individual with PD using the PSER129 specific antibody EP1536Y. As expected^18^, ABC-BAR-PSER129 amplification resulted in a substantial signal enhancement in tissues when compared to standard IHC protocol (Figure 2A). In agreement with what has been reported previously with other low abundance antigens^19^, DAB staining was readily visible with the naked eye with ABC-BAR-PSER129, while the standard IHC protocol required a microscope to verify staining.

**Figure 2.**
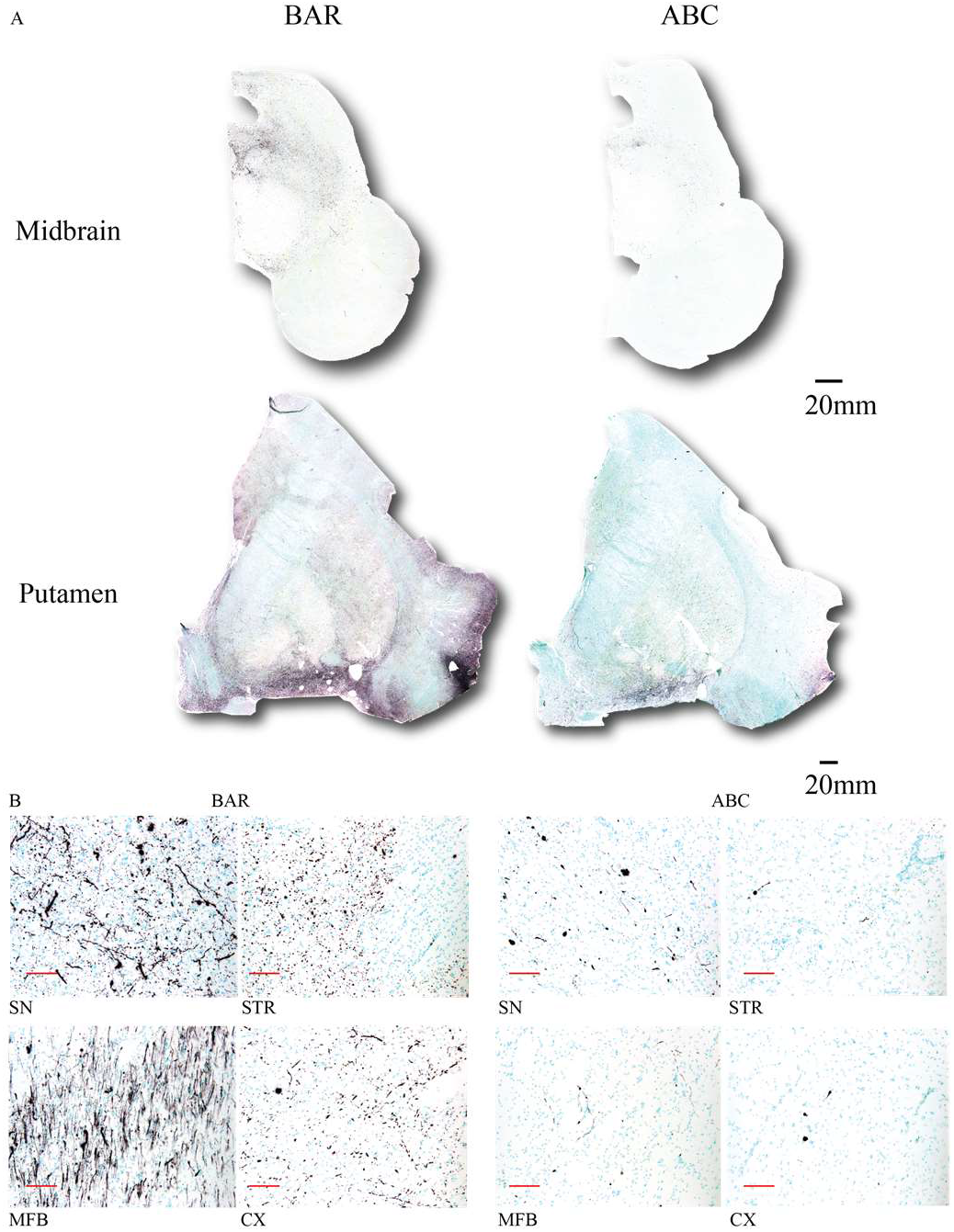
BAR labeling of Lewy pathology in the PD brain. Formalin fixed floating sections from midbrain and adjacent to the anterior commissure of one individual with PD were labeled using the ABC-BAR-PSER129 technique or a standard ABC protocol. Tissues were first probed with the PSER129 antibody EP1536Y and then tyramine signal amplification was used to proximity-label molecules adjacent to the antibody. Labeling was detected using ABC and nickel-DAB chromogen. All slides were counterstained with the nuclear stain methyl green. (A) Whole slide scans at 10X magnification of midbrain and putamen containing sections show distribution and intensity of BAR labeling and using standard ABC detection. (B) 20X images of regions of interest including the substantia nigra (SN), striatum (STR), medial forebrain bundle (MFB), and the cortex (CX). All sections were from the same PD brain sample. Scale bars for B = 100 microns.

The visibility appeared to be a result of both the increase signal intensity and the low background associated with the technique^19^. ABC-BAR-PSER129 amplification heavily labeled cell bodies of the substantia nigra (SN), projections of the medial forebrain bundle (MFB), punctate structures throughout the putamen and some of adjacent cortical regions. In contrast, the standard protocol used on tissue from the same individual, labeled few punctate structures of the putamen, and few projections in forebrain bundle and cortex (Figure 2B). In sum, following ABC-BAR-PSER129 amplification biotin labeling was seen throughout the midbrain, putamen, and cortical regions of PD brain.

Next, we applied the ABC-BAR-PSER129 technique to tissues from both PD and DLB. We found that in all samples tested, ABC-BAR-PSER129 resulted in enhanced pathology detection. Particularly, small punctate structures in the cortex and striatum were more easily observed when compared to standard ABC labeling (Fig 3A). Match cortices from the same individual with DLB, show more apparent small punctate structures for ABC-BAR-PSER129 compared to standard ABC when viewed under high magnification (Figure 3B) .To ensure that the labeling with ABC-BAR-PSER129 technique was indeed specific to Lewy Pathology we then performed several control studies testing the specificity of the ABC-BAR-PSER129 labeling (Fig 3C). PD brain sections digested with proteinase K showed similar ABC-BAR-PSER129 labeling as undigested sections. In contrast, PD brain sections that were incubated with the preabsorbed EP1536Y antibody or processed in the absence of EP1536Y did not show ABC-BAR-PSER129 labeling. Together these results demonstrate that labeling was specific for PSER129 positive Lewy Pathology.

**Figure 3.**
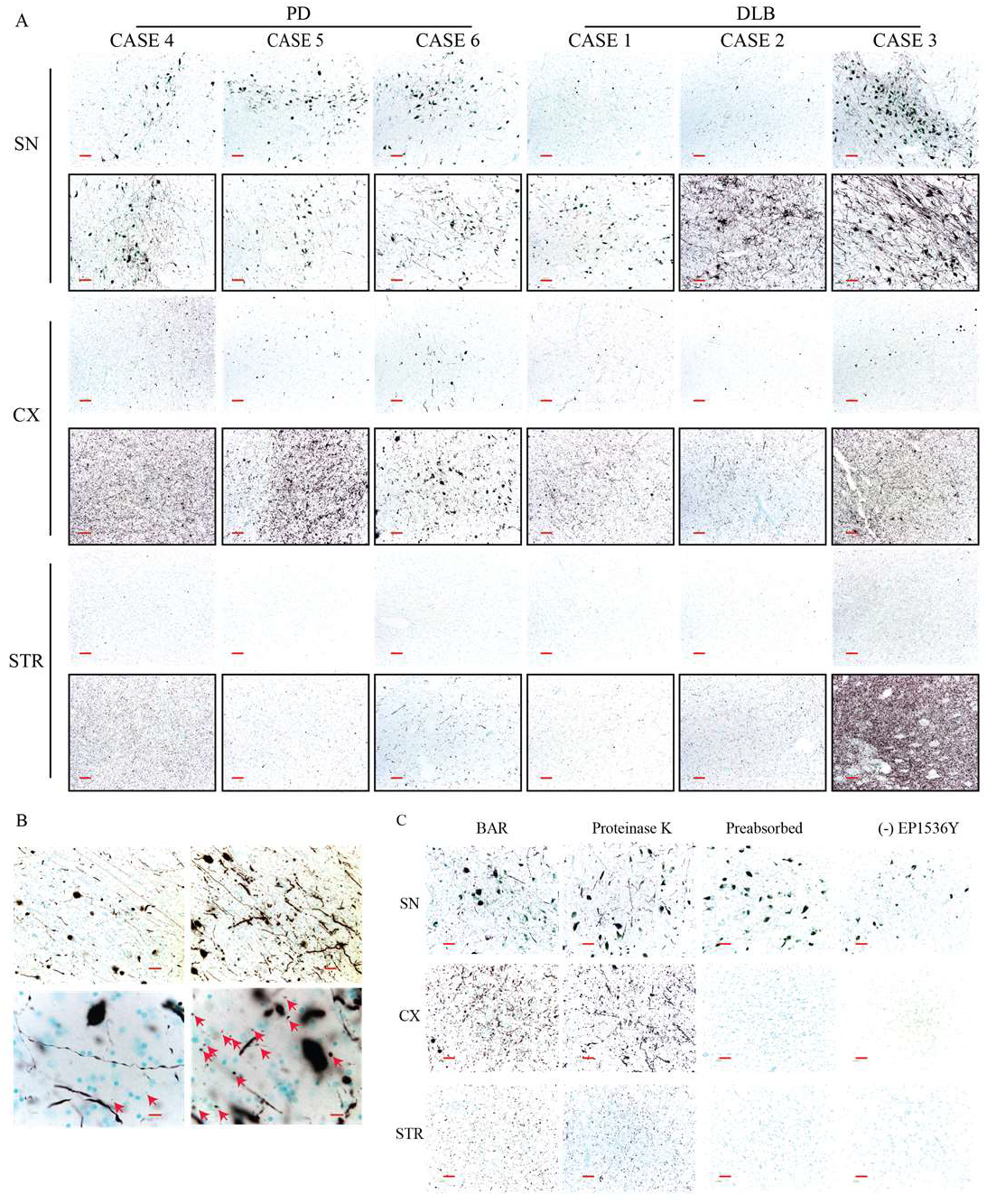
BAR labeled Lewy Pathology in PD and DLB brain. Formalin fixed floating sections from midbrain and putamen of 3 PD and 3 DLB individuals were labeled using the ABC-BAR-PSER129 technique or a standard ABC protocol. (A) 10X images of regions of interest across all individuals. Images with black border are from tissues processed with ABC-BAR-PSER129. Images without a border are from tissues processed with a standard ABC protocol. Labeling was detected using ABC and nickel-DAB chromogen. All slides were counterstained with the nuclear stain methyl green. (B) 10X (top panels) and 100X images (bottom panels) from cortical region baring Lewy Pathology processed with ABC-BAR-PSER129. Red arrows denote small punctate pathology. Scale bar top panels = 50 microns, scale bar bottom panel = 10 microns. Scale bar bottom panels = 10 microns (C) Specificity controls were conducted for the ABC-BAR-PSER129 technique. Sections from individual “case 1” were processed as before using ABC-BAR-PSER129 (BAR). One set of sections were processed with ABC-BAR-PSER129 following 10 min digestion with 20 μg/mL proteinase K (PK). Sections processed with ABC-BAR-PSER129 using antibody EP1536Y preabsorbed with a per129 peptide fragment (PA). Sections processed with ABC-BAR-PSER129 in the absence antibody EP1536Y (-Ab). SN= Substantia Nigra, Medial Forebrain Bundle = MFB, CX = Cortex, STR = Striatum. Scale bar = 50 microns.

To determine whether we could isolate the BAR-labeled pathology from tissue, we then reversed crosslinking and solubilized cross-linked samples using a previous protocol with modifications^20^. Prior to cross-link reversal tissue was washed with SDS and 2-Mercapitoethanol to remove contaminating primary antibody. Following heat mediated crosslink reversal very little pellet was observed after centrifugation (i.e. 22,000 × g for 30 min), suggesting the tissue was effectively solubilized. Following streptavidin pulldown of the labeled midbrain and putamen sample, the lysate contained little to no detectable biotin while the pulldown sample contained abundant biotinylated proteins, at a diverse range of molecular weights (Fig 4A). We then probed the samples with two alpha-synuclein antibodies reactive towards different epitopes, BDSYN1 (aa. 91-99; BD Biosciences) and MJFR1 (a.a. 118-123; Abcam). With both antibodies, we observed predominately high molecular weight species over 75 kDa in the captured sample. Alpha-synuclein that was not captured (i.e. flow through) consists of a mixture of apparent monomer at 15 kDa and high molecular weight species greater than 37 kDa. Next, we probed for PSER129 using the EP1536Y antibody and found high molecular weight species predominantly in the captured sample. Next, we probed for ubiquitin conjugates and observed several bands approximately at 20-25 kDa, consistent with ubiquitinated alpha-synuclein isolated from Lewy pathology^21^. Together, results demonstrated the ABC-BAR-PSER129 captured proteins were immunoreactive for known Lewy Pathology markers.

**Figure 4.**
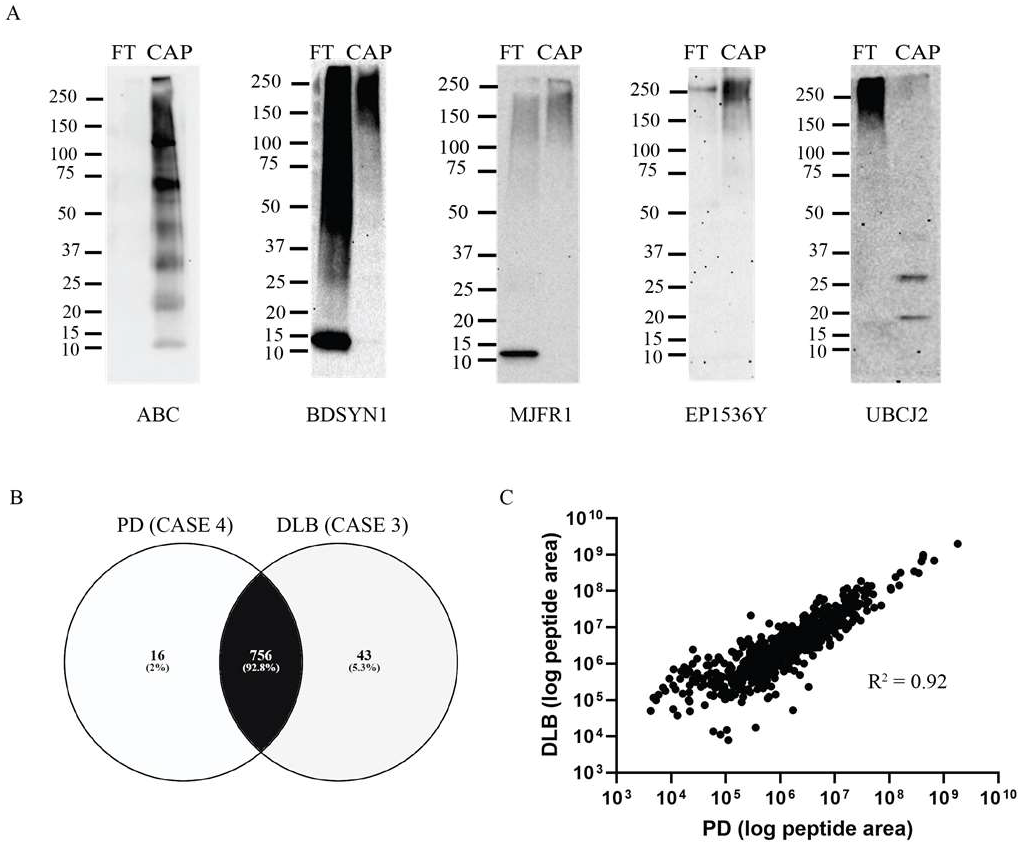
Isolation and mass spectrometry of BAR labeled Lewy Pathology. ABC-BAR-PSER129 was conducted on a single formalin fixed floating section from the putamen and midbrain. Tissues from 2 individuals were processed, “case 4” and “case 3” (See table 1). These tissues were selected because they showed similar distribution and amount of Lewy pathology. Following ABC-BAR-PSER129, residual antibody-ABC complexes were stripped from the tissue. Crosslinks were then reversed and proteins extracted at high temperatures. Extracted biotin labeled proteins were captured using magnetic streptavidin beads. (A) Western blot analysis of captured proteins (CAP) and flow through from bead purification (FT). Blots were then probed with ABC to detect biotinylated proteins in the samples (ABC). Blots were also probed with two antibodies against alpha-synuclein (BDSYN1 and MJFR1). Blots were also probed with anti-pser129 alpha-synuclein antibody (EP1536Y) and the anti-ubiquitin conjugate antibody (UBCJ2). Molecular weight marker has been denoted for each blot. Venn-diagram depicting the number and percentage total proteins identified in each sample and overlap between samples. (C) Pearson correlation analysis relative abundance of proteins between the two samples. Relative abundance of each peptide was estimated using the sum of the areas for the three most abundant peptides assigned to a given protein accession number.

Next, we further characterized ABC-BAR-PSER129 captured Lewy pathology by mass spectrometry. To do this, the midbrain and putamen of one PD and one DLB case were processed with the ABC-BAR-PSER129 protocol, and the captured proteins were digested and identified by mass spectrometry. In total, we identified 815 proteins, with 16 being unique to PD sections and 43 being unique to the DLB sections (Figure 4B; See supplementary file 3 for complete output). The majority of proteins identified (756 proteins or 92.8%) were common between both of the independently processed samples from different individuals. Because proteins identified were highly homologous between the two samples, we wanted to determine if the relative abundance of identified proteins were also similar. To do this we conducted a correlation analysis comparing the two samples using the total area for the top three unique peptides identified for each protein as a measure of relative abundance. We found a significant correlation between the samples (R2=0.92, p<0.0001), demonstrating that the measured relative abundance of proteins was highly similar between the two samples (Figure 4C). Consistent with successful PSER129 purification, alpha-synuclein had 76% percent protein sequence coverage. Together, results showed that mass spectrometry could be used for the identification of proteins from ABC-BAR-PSER129 captured fraction.

Lastly, we investigated whether the proteomic signature of the captured fraction resembled Lewy Pathology. To determine this we conducted a pathway enrichment analysis on the 815 proteins identified, using available mass spectrometry derived brain proteome data^15^ as background (See supplementary file 5 for complete protein list) to avoid artificially inflated enrichment scores (i.e. false positives). Results showed proteins identified were enriched for many annotated processes and pathways (Figure 5A; See supplementary file 1 for complete output). Of particular interest, the most enriched KEGG pathway was Parkinson’s disease (adj. p-value = 7.30 × 10^−32^) and the most enriched for GO cellular compartment was vesicles (adj. p-value = 1.87 × 10^−79^) (Figure 5B). Indeed, the top 6 enriched GO cellular compartments involved extracellular vesicles including extracellular exosomes (adj. p-value = 5.32 × 10^−74^). Several other neurodegenerative disorders appeared in the top 10 KEGG pathways including Prion disease (adj. p-value = 1.15 × 10^−25^), Huntington disease (adj. p-value = 2.75 × 10^−23^), Alzheimer’s disease (adj. p-value = 1.58 × 10^−21^), and Amyotrophic lateral sclerosis (adj. p-value = 1.93 × 10^−15^). Subsequent clustering of pathway enrichment analysis was used to provide a visual overview of cellular pathways and processes associated with the ABC-BAR-PSER129 captured Lewy pathology. The clustering analysis allows major themes to be elucidated, as there is much overlap between annotated pathways. Results showed several apparent major clusters including vesicle trafficking, synaptic neurotransmission, mitochondrial respiration, and antigen presentation amongst others (Figure 5C). ABC-BAR-PSER129 captured Lewy pathology was enriched for cerebral cortex neuropil (adj. p-value = 3.94 × 10^−59^) and substantia nigra development (adj. p-value = 2.8 × 10^−7^), supporting the hypothesis that Lewy pathology is most prevalent in fibers of dopamine neurons. To determine whether Lewy pathology was statistically not associated with a pathway (i.e. underrepresented) we conducted a pathway exclusion analysis with gprofiler. Results showed that particularly transcriptional processes of the nucleus were underrepresented in the ABC-BAR-PSER129 captured Lewy pathology (See supplementary file 2 for complete output), suggesting that processes in the nucleus are not in direct association with synucleinopathy.

**Figure 5.**
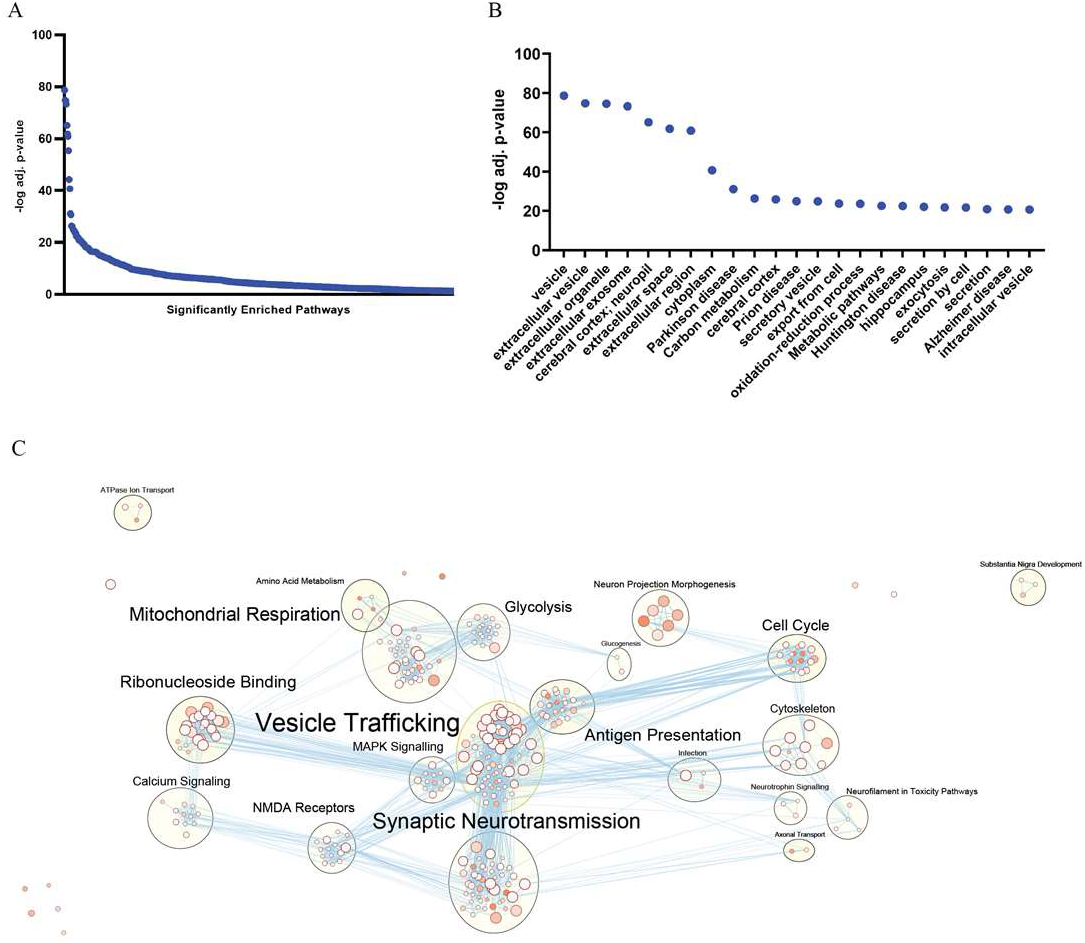
Pathway enrichment analysis of Lewy pathology from the PD and DLB brain. Proteins identified from the ABC-BAR-PSER129 labeled PD and DLB brain were analyzed using gProfiler. Adj. p-values for all significantly enriched pathways. (B) Pathways enriched with a cutoff value of −log10 adj. p-value > 20. (C) Significantly enriched pathways were visualized using Cityscape with Enrichmentmap (q<0.001). Generalized terms have been manually annotated over each cluster of nodes.

## Discussion

Lewy pathology consists of alpha-synuclein along with numerous other molecules^6,22,23^. Understanding the molecular makeup of Lewy pathology is critical for understanding disease. Organelles such as mitochondria, lysosomes, etc. are associated with the pathology and this protocol may isolate those pathology “microdomains” for characterization, and further the understanding of Lewy pathology formation. Here we found that BAR-technique with antibody EP1536Y effectively labeled pathology burdened cellular processes of the midbrain, putamen, and cortex of the PD and DLB brain. The labeling protocol allowed visualization of small punctate structures that were otherwise difficult to detect with standard IHC methods. These structures were specific for PSER129, and in several diseased brains, they represented the dominant pathological structure. Mass spectrometry analysis of these labeled structures strongly suggested they were extracellular vesicles.

There are several major advantages of the BAR-technique. First, this protocol allows for the molecular characterization of Lewy pathology in primary human tissues. This could lead to a deeper understanding of how animal models bearing Lewy pathology represent or do not represent human diseases. Furthermore, it could allow a more detailed understanding of the synucleinopathy disease process across a range of comorbid neurodegenerative diseases. Second, this protocol requires a single tissue section and typical laboratory reagents, making the routine processing many samples feasible. Previous enrichment strategies for Lewy pathology usually required a mass of fresh tissue and extensive protocol optimization; even then, the enriched fractions were difficult to define and conduct for many samples. A third advantage is protein-protein interactions can be studied in the insoluble material of Lewy pathology. A fourth advantage is the potential to understand “pathological domains” in diseased cells. BAR labels molecules based on proximity, and therefore Lewy pathology and its cellular compartment will be labeled, giving not only insight into the molecular makeup of Lewy pathology but also information about the diseased cell, respectively. The last major advantage of the BAR technique is it provides information about the core pathological process for synucleinopathies, while other approaches, for example bulk tissue RNA-seq, measure ancillary cellular processes.

Pathway enrichment analysis was particularly useful for analyzing data produced from ABC-BAR-PSER129. The most significant KEGG pathway was “Parkinson’s Disease.” In total, 12% of identified proteins were from the KEGG pathway “Parkinson’s Disease.” Intriguingly, exosomes were one of the most significantly enriched for GO cellular compartment, and vesicles/extracellular vesicles were also top-hits, providing good evidence that Lewy pathology is found in exosomes in the human diseased brain. This finding has important implications for the hypothesized prion-like spread of Lewy pathology, as the mechanism of intercellular spread remains unclear, and our data suggests Lewy pathology may be primarily spreading via exosomes^24^. If Lewy pathology occurs in extracellular exosomes, it may be possible to target these structures to quell the spread. Adjacent to vesicular trafficking, MHCII antigen presentation and infections may play a component in the disease process. Intuitively, endosomes containing the disease peptide (alpha-synuclein) are exported to the cell surface to trigger an MHII mediated immune response. In support of this interpretation endosomal and secretory granule pathways were common pathways enriched in the vesicular trafficking cluster. Proteins identified around Lewy pathology were uniform between DLB and PD samples supporting a common underlying mechanistic process for both diseases. Applying the ABC-BAR-PSER129 technique to more cases will be important to understand both how cellular pathways are unique to specific synucleinopathies and to determine possible molecular heterogeneity between synucleinopathies.

Several important considerations should be made when using this technique. Only well-characterized primary antibodies used under optimal conditions should be adopted for this protocol. This consideration was our main justification for using antibody EP1536Y to label Lewy pathology, despite many “marker” antibodies being available. Second, the antibody-stripping step included here can be omitted for some applications, such as mass spectrometry, as this harsh method likely also extracts some proteins from the primary tissue. Removing unwanted antibodies and complexes (i.e. avidin-HRP) prior to extraction has the specific benefit of being amenable to western blot detection but may otherwise be unnecessary. Third, the BAR reaction time can be increased or decreased to label structures less or more stringently, respectively. We used a relatively long reaction time to increase the consistency of labeling that is otherwise hard to achieve with shorter reaction times, and also in an effort to label the cellular compartment in which the Lewy pathology was contained. We did not determine the labeling radius but previous reports have found that molecules within 200-300nm range of the antigen are labeled with biotin^9^, although this estimated labeling radius does not take in to account the size of the antibody-ABC-HRP complex utilized here.

An important observation of these studies was the sensitive labeling of PSER129 positive pathology we were able to achieve in fixed tissue. The intense labeling did not require tissue digestion or other common antigen retrieval protocols and agrees with previous reports using similar detection protocols for low abundance antigens^18^. ABC-BAR-PSER129 allowed for more complete labeling, especially of smaller cellular processes harboring pathology. The best example being punctate structures of the striatum and cortex that were otherwise in several individuals brains unlabeled using standard techniques. Indeed, tyramine signal enhancement (TSA) has been used in neural tracing studies because it can drastically enhance the labeling of fine processes in fixed tissues^25^. Besides the ability for ABC-BAR-PSER129 to simplify the IHC protocol, a benefit of assessing pathology in intact heavily cross-linked samples is avoidance of high background and tissue damage sometimes results from antigen retrieval. Furthermore, the high dilutions (>1:50K) of EP1536Y used would make binding kinetics more favorable, and could account for the high signal to background observed with this labeling strategy^19^. The high level of sensitivity allowed the unambiguous identification of PSER129 positive structures in PD tissues without a microscope; which was not achieved in the same samples processed with typical ABC based protocols. Preadsorption of EP1536Y or excluding EP1536Y from the protocol eliminated the majority of labeling, demonstrating specificity and excluding the possibility of endogenous biotin interference. Interestingly, a few PSER129 positive inclusions were still observed in the striatum (data not shown) after preadsorption of the primary antibody, suggesting that very high dilutions (>1:500K) of EP1536Y likely could effectively label tissues using this protocol.

In conclusion, the technique described here can be used for highly sensitive detection and enrichment of Lewy pathology from formalin fixed brain tissues. Conceptually, we have provided a framework to explore the molecular details of Lewy pathology in primary human tissues and this could instigate a systems based approach for understanding the molecular architecture of Lewy pathology^26^. The modified BAR protocol enriched for proteins within Lewy pathology and possibly adjacent to Lewy pathology (i.e. interactors). Although we explored BAR labeling using a single antibody, the assay could easily be adopted to other high affinity antibodies targeting structures of interest. Future studies using this technique could explore crucial questions about the molecular nature of Lewy pathology in the primary disease tissues that could deepen our understanding of synucleinopathies.

## Supporting information

gprofiler enrichment output

gprofiler underrepresentation output

mass spectrometry output

protein list for BAR enrichment analysis

protein list for enrichment analysis background

## Acknowledgements

We would like to thank the Whitehead Institute Proteomics Facility for conducting the mass spectrometry analysis.

## Supplementary Data

Supplementary file 1 = gprofiler enrichment output

Supplementary file 2 = gprofiler underrepresentation output

Supplementary file 3 = mass spectrometry output

Supplementary file 4 = protein list for BAR enrichment analysis

Supplementary file 5 = protein list for enrichment analysis background

